# The intrinsically disordered C-terminal domain of bacteriophage T4 gp32 amplifies pre-existing conformational heterogeneity at DNA replication junctions

**DOI:** 10.64898/2026.09.18.752737

**Authors:** Lulu Enkhbaatar, Peter H. von Hippel, Andrew H. Marcus

**Affiliations:** Institute of Molecular Biology, University of Oregon, Eugene, OR 97403, USA; Center for Optical, Molecular and Quantum Science, University of Oregon, Eugene, OR 97403, USA; Department of Chemistry and Biochemistry, University of Oregon, Eugene, OR 97403, USA

## Abstract

Selective molecular recognition of dynamic nucleic acid structures is fundamental to DNA replication, yet the physical mechanisms by which intrinsically disordered protein domains achieve this selectivity remain poorly understood. Here, we investigate how the intrinsically disordered C-terminal domain (CTD) of the bacteriophage T4 single-stranded DNA-binding protein gp32 contributes to the recognition of ss–dsDNA replication fork junctions. Using absorbance, circular dichroism, and two-dimensional fluorescence spectroscopy (2DFS) of model DNA replication junctions containing an exciton-coupled (Cy3)_2_ probe, we quantified their average solution structures and conformational heterogeneity. Protein-free DNA junctions exhibit distinct conformational distributions that depend on ssDNA arm lengths and local base sequence, demonstrating that this heterogeneity exists prior to protein binding. Although both wild-type gp32 and the CTD-truncated mutant gp32 I* substantially remodel the average junction structure, only wild-type gp32 preserves and amplifies these differences in conformational heterogeneity. These results support a model in which the intrinsically disordered CTD promotes selective molecular recognition by redistributing populations among pre-existing DNA conformational states rather than by stabilizing a single protein-bound conformation. More broadly, our findings suggest that intrinsically disordered protein domains can recognize dynamic nucleic acid substrates by amplifying conformational heterogeneity already present within their intrinsic conformational landscapes.

## I. Introduction

Faithful DNA replication requires the coordinated activities of numerous proteins as short segments of genomic DNA are sequentially unwound and copied (1). During replication, transient single-stranded (ss)–double-stranded (ds) DNA junctions are the sites of many of the protein-mediated reactions required for DNA synthesis. These ss-dsDNA junctions are themselves highly dynamic, rapidly interconverting among multiple conformations that together constitute the conformational landscape encountered by proteins that regulate DNA replication (2,3).

Selective molecular recognition of dynamic DNA junctions is essential for proteins that coordinate DNA replication. The bacteriophage T4 single-stranded DNA-binding protein gp32 provides a well-characterized model system for investigating this recognition process (4). Following monomer nucleation, gp32 binds cooperatively to ssDNA to form gp32–ssDNA filaments that undergo repeated cycles of assembly and disassembly during DNA synthesis (4,5). A recent study further demonstrated that gp32 preferentially nucleates at primer-template ss– dsDNA junctions containing a single overhanging ssDNA arm with a free 3′ end rather than a free 5′ end (6). These findings revealed that junction recognition is both highly selective and strongly dependent on junction structure.

Insight into this recognition mechanism can be gained by considering the modular domain architecture of gp32. Gp32 consists of a structured ssDNA-binding core domain flanked by intrinsically disordered N- and C-terminal “tail” domains (Fig. 1a) (7). The structured core mediates ssDNA binding, whereas the disordered N-terminal domain stabilizes cooperative interactions between neighboring gp32 monomers (8,9). In contrast, the intrinsically disordered C-terminal domain (CTD) has long been implicated in interactions with other proteins involved in DNA replication (8,10), although whether it also contributes directly to the selective recognition of ss–dsDNA junctions remains unknown.

**Figure 1.**
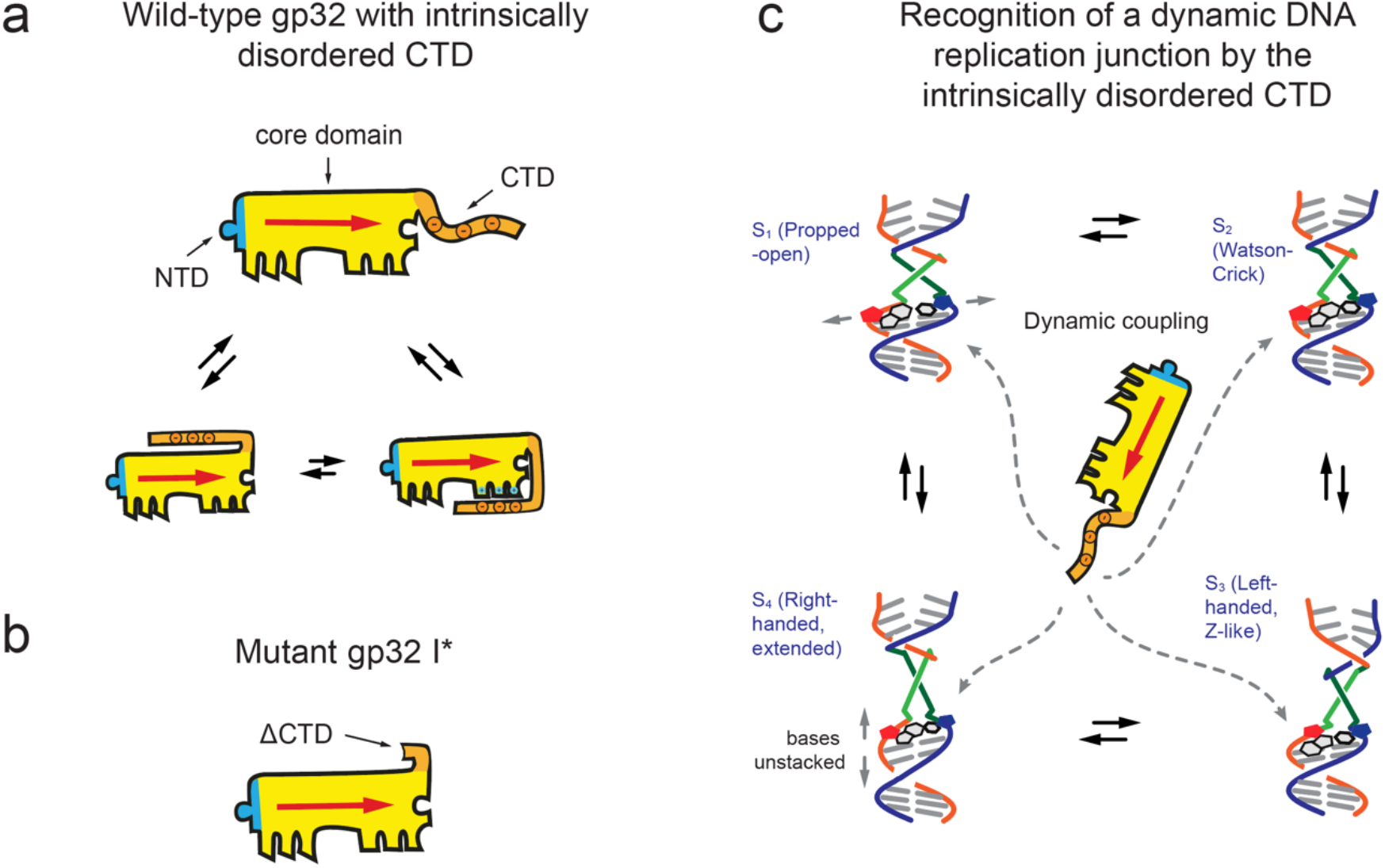
Domain architecture of gp32 and conceptual model for selective recognition of dynamic DNA replication junctions. **(a)** The gp32 monomer consists of an intrinsically disordered N-terminal domain (NTD; residues 1-21), a structured core ssDNA-binding domain (residues 22-253), and an intrinsically disordered C-terminal domain (CTD; residues 254-301). A gp32 monomer, when bound to ssDNA, occupies seven nucleotides (6). The CTD is shown schematically sampling multiple conformations, reflecting its intrinsic structural disorder and dynamic interactions. **(b)** The gp32 I* mutant comprises residues 1-253 and therefore lacks the 48-residue CTD while retaining the structured core DNA-binding domain and NTD. Comparison of wild-type gp32 and gp32 I* enables the functional contributions of the CTD to be isolated. **(c)** Previous studies showed that protein-free DNA replication junctions containing an exciton-coupled (Cy3)_2_ probe sample four representative conformational macrostates (S_1_–S_4_) that rapidly interconvert. The green and gray bars represent the two Cy3 chromophores of the exciton-coupled probe. The present study investigates whether the intrinsically disordered CTD promotes selective molecular recognition by dynamically modulating this pre-existing conformational ensemble rather than by stabilizing a single DNA conformation. Panel a adapted from (6), panel c adapted from (3).

Intrinsically disordered protein regions are increasingly recognized as important contributors to molecular recognition, where their conformational flexibility can facilitate selective interactions with protein and nucleic acid partners (11,12). However, much less is understood about whether disordered protein domains can promote selective recognition by modulating the pre-existing conformational landscapes of their nucleic acid substrates. Current models attribute several dynamic regulatory functions to the intrinsically disordered CTD, including competition with ssDNA for access to the ssDNA-binding cleft, recruitment of regulatory proteins, and transient interactions with DNA that facilitate efficient cycles of gp32 binding and dissociation (5,13). Together, these observations suggest that the intrinsically disordered CTD regulates gp32 through dynamic interactions with DNA, raising the possibility that selective molecular recognition is achieved by modulating the conformational landscape of ss–dsDNA junctions rather than by stabilizing a single DNA conformation. The gp32 I* mutant (Fig. 1b), which lacks the 48-residue CTD, retains tight, cooperative binding to ssDNA and helix-destabilizing activity while exhibiting altered interactions with replication proteins (10). Gp32 I* therefore provides a useful tool for isolating the functional contributions of the CTD.

### Alt text

Three-panel schematic. Panel a shows wild-type gp32 bound to ssDNA, with an N-terminal domain, structured ssDNA-binding core, and intrinsically disordered C-terminal domain depicted in multiple conformations. Panel b shows the CTD-truncated gp32 I* protein, which retains the N-terminal domain and DNA-binding core. Panel c shows four interconverting conformational macrostates (S_1_–S_4_) of a DNA replication junction containing two Cy3 chromophores near the ss–dsDNA junction, together with the gp32 CTD dynamically sampling interactions with the junction conformational ensemble.

To investigate the mechanism of selective junction recognition illustrated schematically in Fig. 1c, we characterized the conformational heterogeneity of a series of model DNA replication junctions in the absence of protein and in separate complexes with wild-type gp32 and the CTD-truncated gp32 I*\** mutant. We employed DNA constructs in which an exciton-coupled Cy3 dimer was incorporated into the sugar-phosphate backbones near the ss–dsDNA junction (14). Because the optical transitions of the exciton-coupled dimer are exquisitely sensitive to the local DNA conformation, our absorbance, circular dichroism (CD), and two-dimensional fluorescence spectroscopy (2DFS) data provide complementary measures of the average solution structure and conformational heterogeneity of the junction (15,16). 2DFS is particularly sensitive to conformational disorder within the DNA immediately surrounding the fluorescent probe (14,16,17). Together, these measurements provide a direct test of whether the intrinsically disordered CTD contributes to selective junction recognition by modulating pre-existing DNA conformational heterogeneity.

## II. Materials and Methods

### DNA constructs and protein preparation

The (Cy3)_2_-labeled ss–dsDNA replication junction constructs were prepared by annealing complementary oligonucleotides obtained from Integrated DNA Technologies (IDT, Coralville, IA). Constructs differed in the lengths of their 3′ and 5′ ssDNA arms while maintaining a common duplex region and (Cy3)_2_ probe position. An additional construct in which A and T were exchanged between the two strands was prepared to assess the influence of local base sequence. Complete oligonucleotide sequences are provided in Supplementary Table S1. Wild-type gp32 was expressed and purified as described previously (18). The gp32 I* expression plasmid was provided by Richard Karpel, and the protein was expressed and purified using the same procedure. Purified proteins were dialyzed into the experimental buffer, and protein concentrations were determined spectrophotometrically at 280 nm using a molar extinction coefficient of 37,000 M^-1^ cm^-1^ for both wild-type gp32 and gp32 I* (19). All samples were prepared in buffer containing 100 mM NaCl, 6 mM MgCl_2_, and 10 mM Tris (pH 7.5). Additional details about sample preparation are provided in the Supplementary Information.

### Absorbance and circular dichroism (CD) spectroscopy

Absorbance and CD spectra of 1 μM DNA samples were measured using a JASCO J-720 spectropolarimeter at 21 °C. Saturating stoichiometric concentrations of wild-type gp32 and gp32 I* were determined independently for each DNA construct by protein titration until no further change in the CD spectrum was observed. Saturation was achieved at DNA:protein molar ratios of 1:3 for the 3’9A/5’2T and 3’2A/5’9T constructs, 1:4 for the 3’9A/5’9T reference construct, and 1:5 for the 3’22A/5’9T and 3’9A/5’22T constructs. Titrations were performed in duplicate. Additional experimental details are provided in the Supplementary Information.

### Two-dimensional fluorescence spectroscopy (2DFS)

2DFS measurements were performed using phase-modulated fluorescence-detected methods described previously (16,17,20). DNA samples (1 μM) were continuously circulated through the optical cuvette and connecting tubing using a peristaltic pump during data acquisition. Rephasing (RP) and nonrephasing (NRP) signals were acquired as functions of the coherence time delays with the population time set to *T* = 0 and Fourier transformed to obtain the corresponding two-dimensional frequency-domain spectra. The frequency axes were referenced using optical reference signals as described previously (20). For each DNA construct, two measurements were initially acquired in the absence of protein to assess reproducibility. Protein was then added at the appropriate saturating stoichiometric concentration, and the protein–DNA complex was equilibrated for 10 min at room temperature before a single 2DFS measurement was acquired for each protein-DNA condition (*n* = 1). The excitation spectrum, optical power, and temporal overlap of the excitation pulses were monitored throughout data acquisition and recalibrated as necessary. Additional experimental details are provided in the Supplementary Information.

### Spectral modeling and analysis

Absorbance and CD spectra were analyzed using a Holstein– Frenkel Hamiltonian and transition-charge description of the excitonic coupling between the two Cy3 chromophores, as described previously (15,17,21). Simultaneous optimization of the absorbance and CD spectra was used to determine the ensemble-averaged structural parameters of the (Cy3)_2_ probe. These structural parameters were subsequently held fixed while the RP and NRP 2DFS signals were analyzed simultaneously to determine the homogeneous and inhomogeneous lineshape parameters, *Γ*_*H*_ and *σ*_*I*_, respectively. Parameter uncertainties were determined as described previously (17), with uncertainty bounds defined for the present analysis by parameter variations satisfying 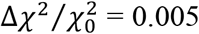, corresponding to a 0.5% increase relative to the optimized 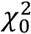 value.

### Use of Artificial Intelligence

ChatGPT (OpenAI) was used during the preparation of this manuscript to assist with language editing and refinement of the presentation. All AI-assisted edits were reviewed and approved by the authors.

## III. Results

### Protein-free DNA junctions exhibit architecture-dependent conformational heterogeneity

To investigate the role of the gp32 CTD in selective junction recognition, we employed a series of model DNA replication junctions containing an exciton-coupled (Cy3)_2_ probe positioned adjacent to the ss–dsDNA junction (Fig. 2). Relative to the common 3′9A/5′9T reference architecture, the constructs systematically vary the length of either the 3′ or the 5′ ssDNA arm while maintaining a common duplex region and probe position. A final construct, in which A and T were exchanged between the two strands, was included to assess the influence of local base sequence at the model DNA fork junction.

**Figure 2.**
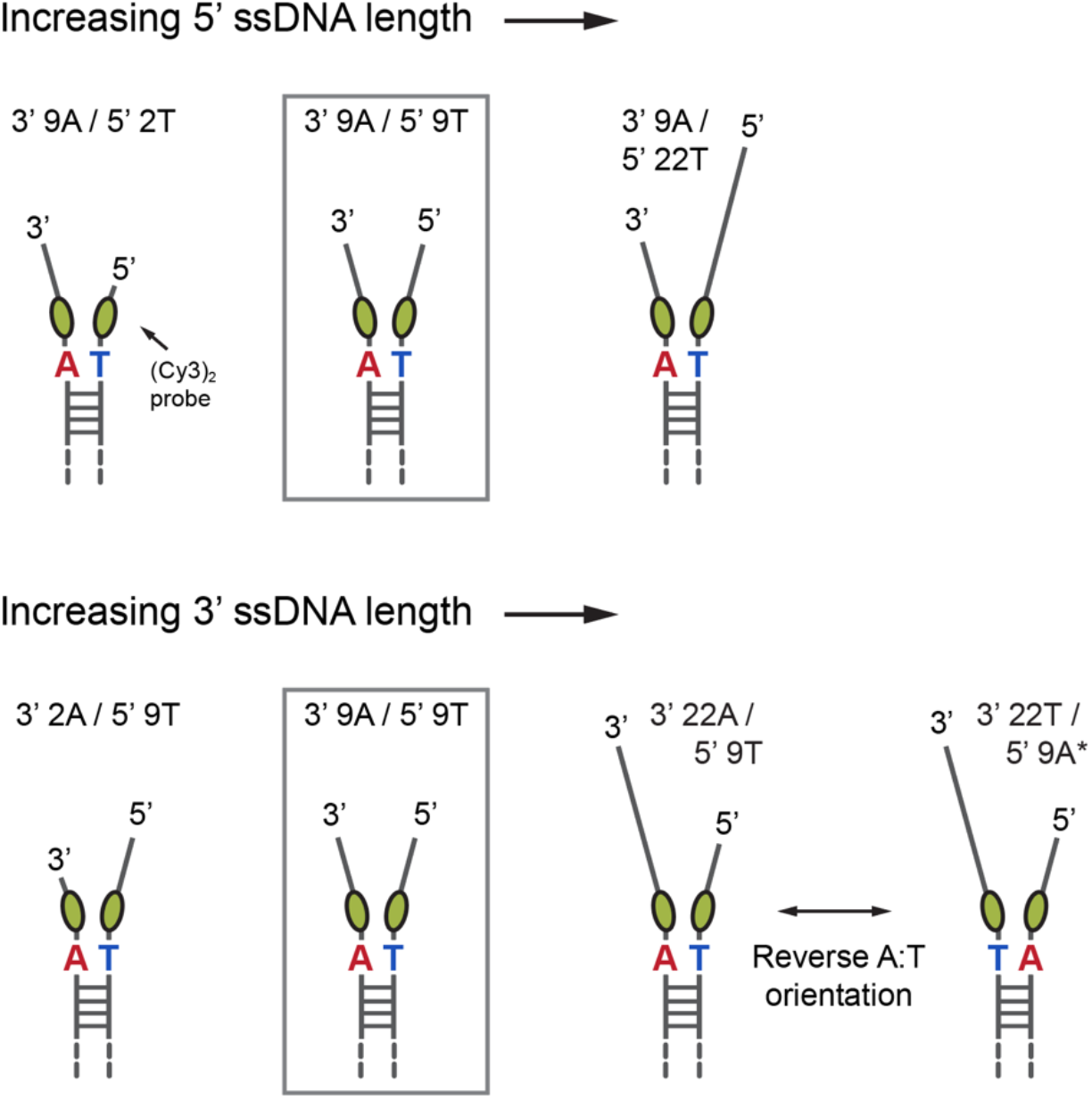
Model DNA replication junctions used in this study. All constructs contain an exciton-coupled (Cy3)_2_ probe positioned at the −1 base pair adjacent to the ss–dsDNA junction. The lengths of the 5′ or 3′ ssDNA arms were varied systematically while maintaining a common duplex region and probe position. The 3′9A/5′9T construct serves as the reference architecture for both series. An additional construct in which A and T were exchanged between the two strands (3′22T/5′9A*) was used to assess the effects of local base sequence at the model DNA fork junction. Complete nucleotide sequences are provided in Supplementary Table S1.

#### Alt text

Schematic series of six DNA replication fork junction constructs containing two Cy3 chromophores at the −1 base pair adjacent to the ss–dsDNA junction. The constructs share a common duplex region but differ in the lengths of their two ssDNA arms, with 3′ and 5′ arms of 2, 9, or 22 nucleotides arranged around the 3′9A/5′9T reference architecture. The final construct, 3′22T/5′9A*, has the same arm-length architecture as 3′22A/5′9T but with A and T exchanged between the two strands.

The exciton-coupled (Cy3)_2_ probe provides a sensitive reporter of the local DNA conformation at the ss–dsDNA junction. Excitonic interactions between the two Cy3 chromophores depend strongly on their relative separation and orientation, allowing changes in the local junction structure to be monitored spectroscopically. Absorbance and circular dichroism (CD) measurements report the average local structure of the probe-labeled junction, whereas two-dimensional fluorescence spectroscopy (2DFS) is additionally sensitive to conformational disorder within the ensemble of structures sampled in solution (15-17). Together, these measurements permit us to quantify both the average junction structure and its conformational heterogeneity.

Before examining the effects of gp32 binding, we first characterized the intrinsic conformational landscape of each protein-free DNA replication junction using absorbance, CD, and 2DFS measurements. Whereas absorbance and CD report the average local junction structure, the 2DFS-derived inhomogeneous linewidth parameter, *σ*_*I*_, provides a quantitative measure of the conformational disorder within the probe-labeled junction (16,17) (Fig. 3a). The absorbance and CD spectra were analyzed globally using a Holstein–Frenkel Hamiltonian to determine the average local geometry of the (Cy3)_2_ probe (15). Representative global analysis of the 3′9A/5′9T reference junction is shown in Supplementary Fig. S1, with experimental and optimized spectra for the complete series of DNA constructs and protein complexes shown in Supplementary Figs. S4 and S5 and the optimized structural and lineshape parameters listed in Supplementary Table S2. Although the average structural parameters vary among constructs, the most pronounced architecture-dependent differences occur in the inhomogeneous linewidth, *σ*_*I*_, indicating substantial differences in conformational heterogeneity, as discussed below.

**Figure 3.**
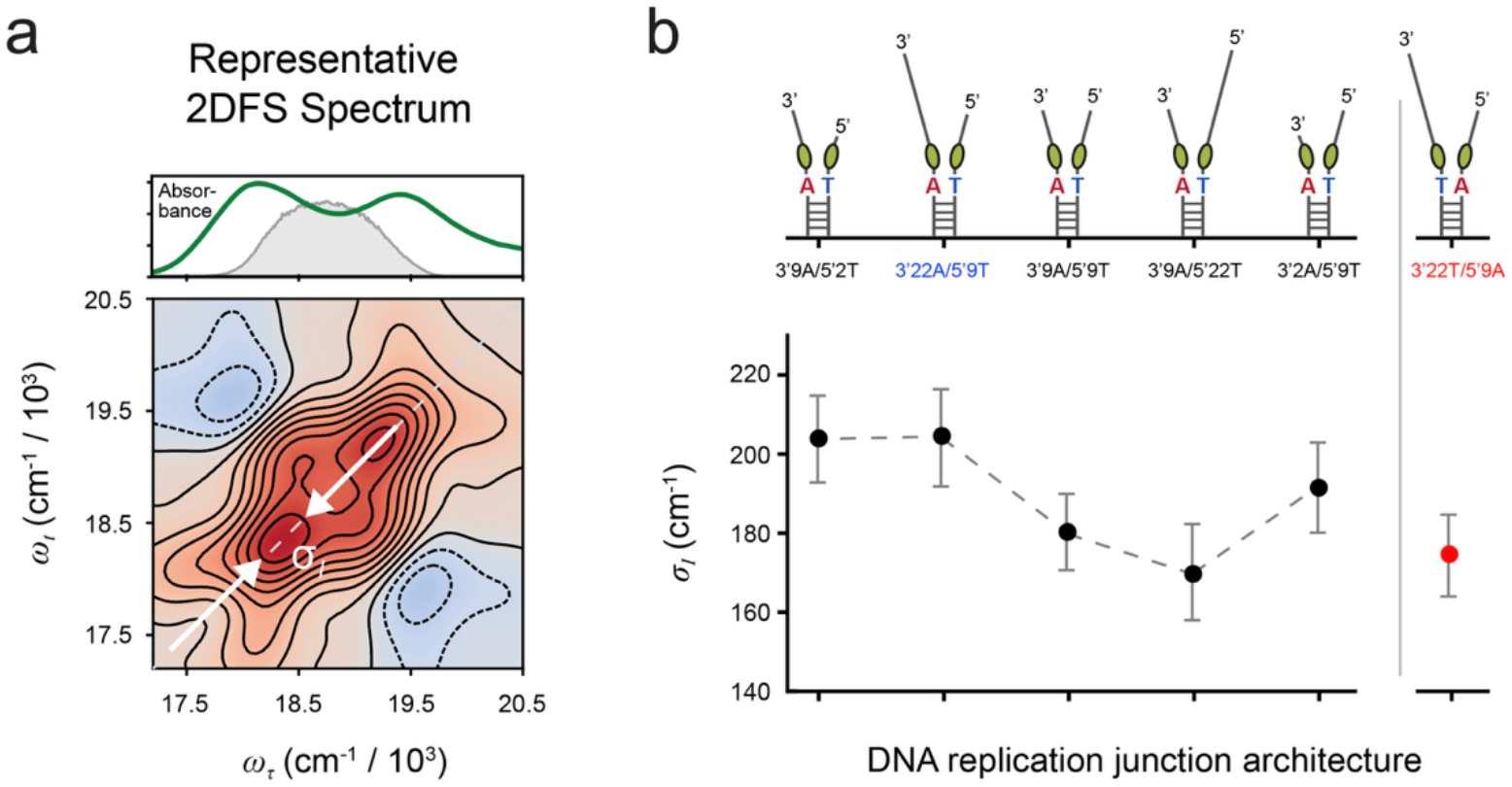
DNA junction architecture strongly influences conformational disorder. **(a)** Representative two-dimensional fluorescence spectrum (2DFS) of an exciton-coupled (Cy3)_2_ probe incorporated at the −1 position of an ss–dsDNA junction. The inhomogeneous linewidth parameter, *σ*_*I*_, obtained from analysis of the 2DFS spectrum provides a quantitative measure of conformational disorder within the local DNA environment surrounding the probe. The upper panel compares the linear absorbance spectrum of the (Cy3)_2_ probe (green) with the excitation laser spectrum (gray). **(b)** Values of *σ*_*I*_ extracted for the six protein-free DNA replication junctions shown in Fig. 2. Constructs are arranged from left to right in order of decreasing 3′/5′ ssDNA arm-length ratio. The final construct, 3′22T/5′9A*, differs from the preceding 3′22A/5′9T junction by exchange of A and T between the two strands adjacent to the (Cy3)_2_ probe and serves as a control for local sequence effects. Error bars represent uncertainties obtained from the global fitting analysis.

#### Alt text

Two-panel figure showing 2DFS measurement of conformational disorder in protein-free DNA replication junctions. Panel a shows a representative two-dimensional fluorescence spectrum of the exciton-coupled (Cy3)_2_ probe, with diagonal and off-diagonal spectral features; above it are the linear absorbance spectrum of the probe and the excitation laser spectrum. Panel b plots the fitted inhomogeneous linewidth parameter, *σ*_*I*_, for six DNA junction architectures arranged in order of decreasing 3′/5′ ssDNA arm-length ratio. The values vary substantially and non-monotonically among the junction architectures, demonstrating architecture-dependent conformational disorder; the 3′22T/5′9A* sequence-exchange control also differs from the corresponding 3′22A/5′9T construct. Error bars show uncertainties from the global fitting analysis.

The conformational heterogeneity of the protein-free DNA replication junctions depends strongly on junction architecture (Fig. 3b). The extracted values of *σ*_*I*_ vary substantially among the six constructs, demonstrating that the junctions sample distinct conformational ensembles even in the absence of protein. *σ*_*I*_ does not vary monotonically with the ratio of the 3′ and 5′ ssDNA arm lengths, indicating that conformational disorder is not determined by this geometric parameter alone. Constructs containing a short two-nucleotide overhang exhibit substantially larger values of *σ*_*I*_ than expected from the overall trend, suggesting that severe shortening of either single-stranded arm introduces an additional source of conformational disorder. If the short-overhang constructs are excluded from this comparison, *σ*_*I*_ decreases systematically with decreasing 3′/5′ ssDNA arm-length ratio. Exchanging A and T between the two strands immediately adjacent to the (Cy3)_2_ probe substantially reduces *σ*_*I*_, demonstrating that local base sequence also contributes to the conformational heterogeneity of the junction. Taken together, these results establish that protein-free DNA replication junctions possess an intrinsic, architecture-dependent conformational landscape that serves as the substrate for subsequent recognition and binding of gp32.

### Wild-type gp32 and gp32 I* produce distinct changes in average junction structure

Having established that the protein-free junctions possess intrinsic, architecture-dependent conformational landscapes, we next examined how wild-type gp32 and the CTD-truncated gp32 I* mutant alter the average local structure of the DNA junction. Binding of either protein produced pronounced changes in the absorbance and CD spectra of the exciton-coupled (Cy3)_2_ probe. However, the spectral changes induced by wild-type gp32 differed substantially from those induced by gp32 I*, indicating that the two proteins produce distinct changes in the average local junction structure. These effects were qualitatively similar across the five junction architectures examined, although their magnitudes varied among constructs (see Supplementary Figs. S2 and S3).

To characterize these structural changes in detail, we examined the CD and absorbance spectra of the 3′9A/5′9T reference construct (Fig. 4). Both wild-type gp32 and the CTD-truncated mutant gp32 I* produce pronounced changes in the CD spectrum relative to protein-free DNA, demonstrating that both proteins substantially remodel the local junction structure. The corresponding absorbance spectra exhibit complementary changes consistent with altered excitonic interactions between the (Cy3)_2_ chromophores. We quantitatively analyzed the CD and absorbance spectra using a Holstein–Frenkel Hamiltonian to determine the ensemble-averaged structural parameters of the (Cy3)_2_ probe for each DNA construct (Supplementary Table S2). Similar qualitative spectral changes were observed for the remaining four DNA junction architectures (Supplementary Fig. S4).

**Figure 4.**
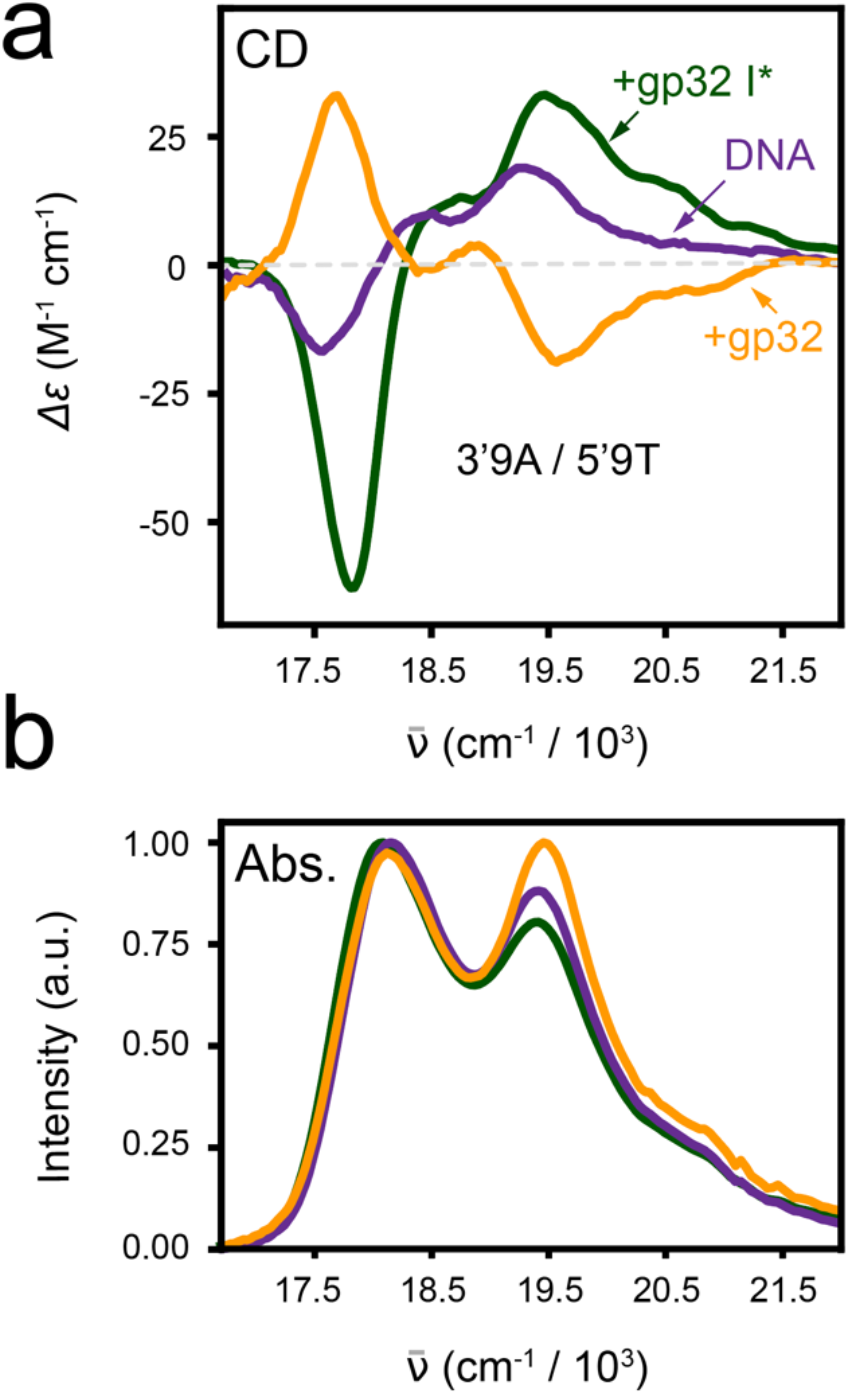
Wild-type gp32 and gp32 I* remodel the average local structure of the DNA replication junction. **(a)** Circular dichroism (CD) spectra of the 3′9A/5′9T reference DNA replication junction in the absence of protein (purple) and following saturation with wild-type gp32 (orange) or the CTD-truncated mutant gp32 I* (green). **(b)** Absorbance spectra normalized to their respective peak absorbance values are shown for the same DNA construct and protein conditions. The CD and absorbance spectra were analyzed simultaneously using a Holstein–Frenkel Hamiltonian to determine the ensemble-averaged structural parameters of the exciton-coupled (Cy3)_2_ probe. Representative and complete spectral fits are shown in Supplementary Figs. S1 and S4.

#### Alt text

Two-panel comparison of CD and absorbance spectra for the 3′9A/5′9T reference DNA replication junction under protein-free, wild-type gp32-bound, and gp32 I*-bound conditions. Panel a shows distinct changes in the CD spectral line shapes upon binding of wild-type gp32 or gp32 I* relative to protein-free DNA, with the two protein-bound spectra also differing from one another. Panel b shows corresponding peak-normalized absorbance spectra, which also exhibit protein-dependent changes in spectral line shape.

The structural analysis indicates that the protein-free DNA junction exhibits a weakly left-handed average structure (*Φ* > 90°), consistent with an ensemble containing both left- and right-handed local conformations. Binding of wild-type gp32 shifts the ensemble toward a pronounced right-handed average structure (*ϕ* < 90°), whereas binding of gp32 I* favors a pronounced left-handed average structure. Thus, although both proteins substantially remodel the average local structure of the DNA replication junction, they produce distinct structural effects. The effects of CTD truncation were particularly revealing for the single-gp32-binding 3′9A/5′2T and 3′2A/5′9T constructs. For the 3′9A/5′2T junction, wild-type gp32 shifts the ensemble toward a right-handed conformation, whereas gp32 I* favors a left-handed conformation; in contrast, both proteins favor a left-handed conformation for the oppositely oriented 3′2A/5′9T junction (Supplementary Fig. S4c, d).

### The gp32 CTD amplifies architecture-dependent conformational heterogeneity

Although the CD and absorbance spectra reveal how gp32 binding alters the ensemble-averaged structure of the DNA replication junction, they provide little information about the breadth of the underlying conformational ensemble. The architecture-dependent disorder observed for the protein-free DNA junctions (Fig. 3) suggests that changes in conformational heterogeneity might constitute an additional consequence of the CTD. To address this question, we analyzed the corresponding two-dimensional fluorescence spectra to determine how wild-type gp32 and gp32 I* differ in their effects on junction conformational heterogeneity.

Among the three representative replication junction architectures examined, protein-free DNA exhibited a systematic decrease in the inhomogeneous linewidth parameter, *σ*_*I*_, as the 3′/5′ ssDNA arm-length ratio decreased (Fig. 5a), consistent with the architecture-dependent conformational disorder established in Fig. 3. Binding of wild-type gp32 preserved this trend but increased the separation between the three junction architectures, indicating that the architecture dependence of conformational disorder is substantially amplified (Fig. 5b). In striking contrast, binding of the CTD-truncated mutant gp32 I* produced nearly identical *σ*_*I*_ values for all three constructs (Fig. 5c), despite maintaining a level of conformational disorder comparable to that observed for the protein-free DNA. The complete optimized structural and lineshape parameters for all constructs and protein complexes are provided in Supplementary Table S2. Thus, removal of the CTD does not eliminate conformational disorder itself; rather, it abolishes its dependence on DNA junction architecture.

**Figure 5.**
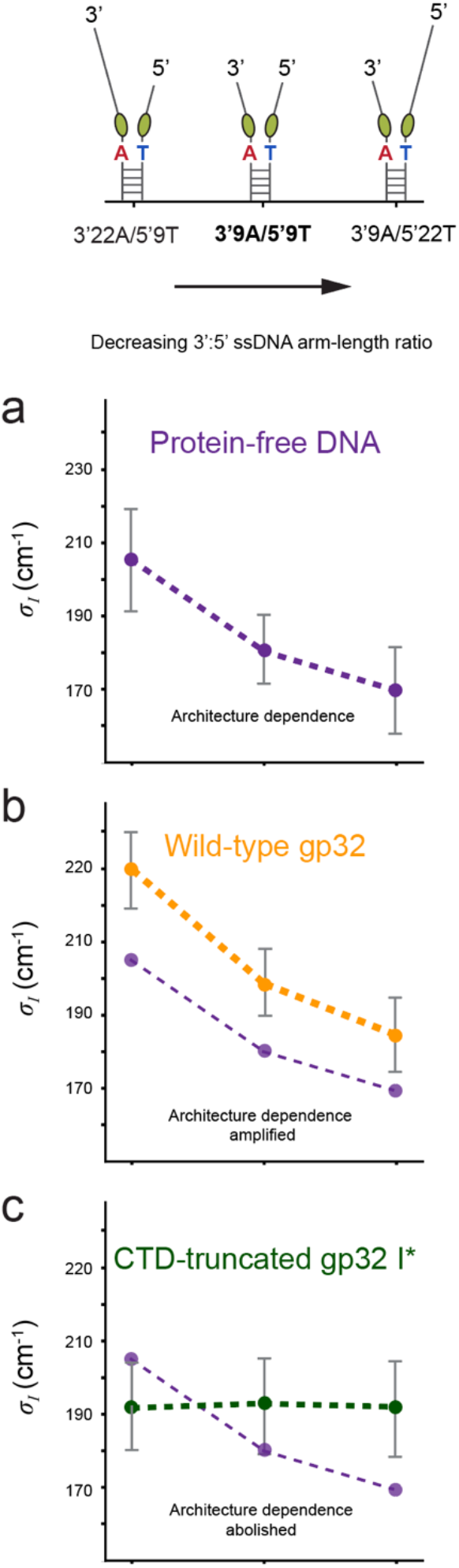
The gp32 CTD amplifies architecture-dependent conformational disorder at DNA replication junctions. Inhomogeneous linewidths (*σ*_*I*_) extracted from 2DFS measurements are shown for three representative DNA replication junction architectures spanning a range of 3′/5′ ssDNA arm-length ratios. The corresponding junction architectures are illustrated above the plots. **(a)** Protein-free DNA exhibits a systematic decrease in *σ*_*I*_ with decreasing 3′/5′ ssDNA arm-length ratio. **(b)** Binding of wild-type gp32 preserves this trend while increasing the differences in *σ*_*I*_ among the three junction architectures. Protein-free DNA values from (a) are reproduced in light purple for direct comparison. **(c)** Binding of the CTD-truncated mutant gp32 I* produces nearly identical *σ*_*I*_ values for all three junction architectures while retaining substantial conformational disorder. Error bars represent uncertainties obtained from the global fitting analysis of the 2DFS spectra.

#### Alt text

Three-panel comparison of the inhomogeneous linewidth parameter, *σ*_*I*_, for three DNA replication junction architectures with decreasing 3′/5′ ssDNA arm-length ratios. Schematics of the three junction architectures are shown above the plots. Panel a shows decreasing *σ*_*I*_ across the three protein-free junctions. Panel b compares the protein-free values with wild-type gp32-bound values, which show larger differences in *σ*_*I*_ among the three architectures. Panel c compares the protein-free values with gp32 I*-bound values, which are similar among the three architectures while remaining substantially above zero.

Taken together, these observations demonstrate that the gp32 CTD does not simply alter the average structure of DNA replication junctions. Rather, it amplifies the intrinsic architecture-dependent conformational disorder already present in protein-free DNA. In contrast, removal of the CTD uncouples conformational disorder from junction architecture while preserving substantial structural heterogeneity. These findings suggest that selective recognition of replication junctions by gp32 arises through modulation of a pre-existing conformational landscape rather than through stabilization of a single preferred DNA conformation.

## IV. Discussion

The present results support a mechanistic model in which selective recognition of DNA replication fork junctions by gp32 arises through dynamic modulation of a pre-existing conformational landscape rather than through stabilization of a single preferred DNA conformation. Protein-free replication junctions populate architecture-dependent conformational ensembles whose breadth is influenced by ssDNA arm lengths and local base sequence. Binding of wild-type gp32 preserves the architecture dependence of this pre-existing conformational heterogeneity while substantially amplifying the differences among junctions. In contrast, removal of the intrinsically disordered C-terminal domain largely abolishes the architecture dependence of the disorder despite producing pronounced changes in the average junction structure. Together, these observations identify the CTD as a key determinant that couples gp32 recognition to the intrinsic conformational landscape of the DNA replication junction.

How can the intrinsically disordered CTD amplify conformational disorder without simply destabilizing the DNA junction? One plausible explanation is that the CTD need not create new DNA conformations but instead redistributes the equilibrium populations of conformational macrostates already sampled by the protein-free junction. Our previous studies demonstrated that replication junctions exist as rapidly interconverting ensembles comprising four representative conformational macrostates (3). We propose that transient interactions involving the CTD redistribute populations among these pre-existing conformational macrostates, thereby amplifying differences among the conformational ensembles while preserving their underlying architecture dependence. This working model is summarized schematically in Fig. 6.

**Figure 6.**
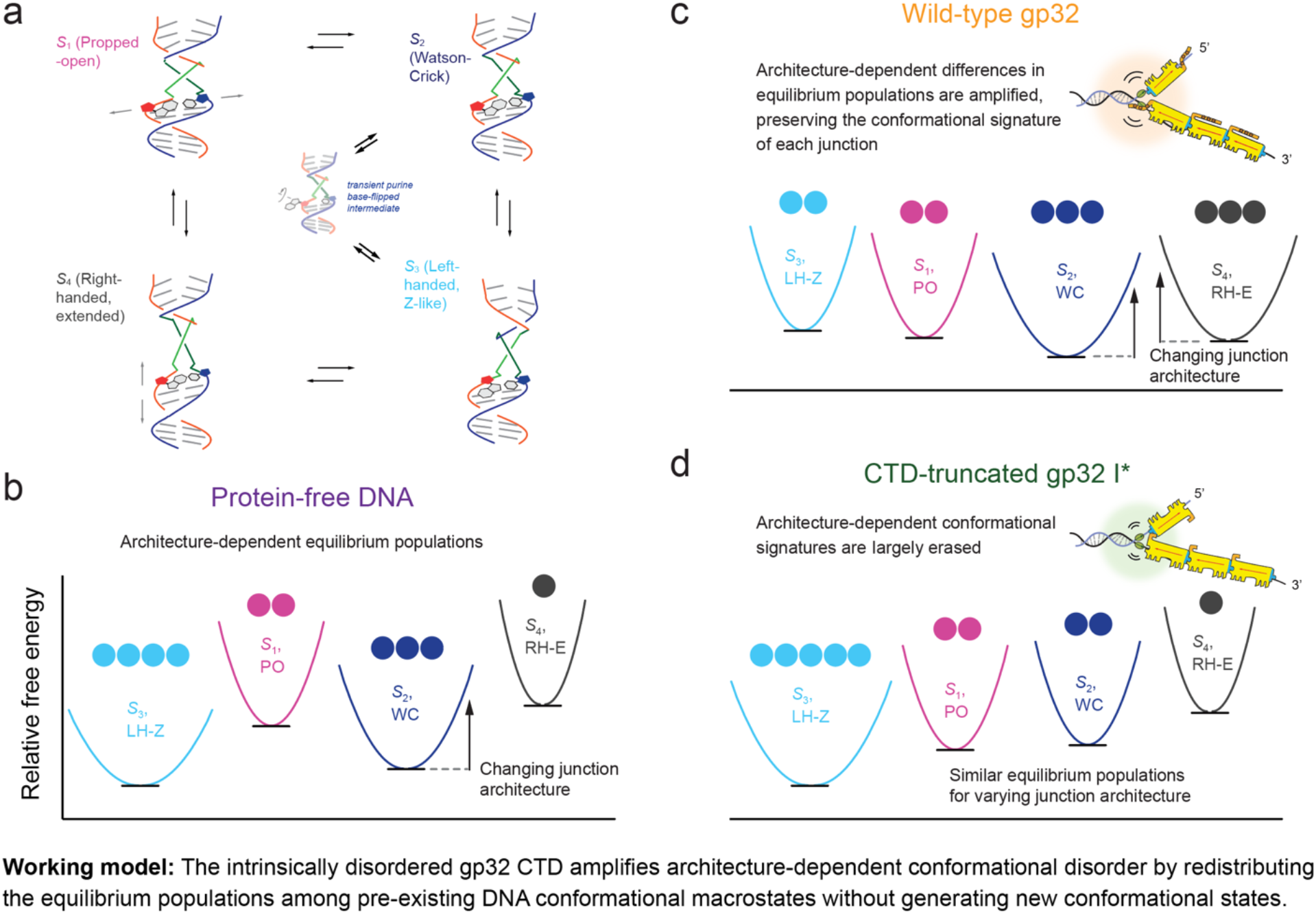
Proposed mechanism by which the gp32 C-terminal domain amplifies architecture-dependent conformational disorder. **(a)** Four conformational macrostates previously identified for protein-free DNA replication junctions (Fig. 1c), together with proposed pathways connecting them. Interconversion between the Watson–Crick (S_2_) and left-handed Z-like (S_3_) conformations is proposed to occur through a transient purine base-flipping intermediate, as suggested previously (3). **(b)** Working model for protein-free DNA replication junctions, in which different junction architectures sample the same set of conformational macrostates but with different equilibrium populations, giving rise to the architecture-dependent conformational heterogeneity observed experimentally (Fig. 3). **(c)** Proposed model for wild-type gp32 binding, in which the intrinsically disordered C-terminal domain (CTD) redistributes populations among pre-existing conformational macrostates, thereby amplifying the dependence of the conformational ensemble on junction architecture (Fig. 5). **(d)** Proposed model for binding of the CTD-truncated mutant gp32 I*, in which the architecture dependence of the conformational ensemble is largely abolished, producing ensembles of similar breadth for different junction architectures (Fig. 5). This working model suggests that the intrinsically disordered gp32 CTD promotes selective molecular recognition by amplifying pre-existing differences among the conformational distributions of DNA replication junctions rather than by inducing a unique DNA structure. Panel a adapted from (3).

The conformational macrostates identified in our previous studies of protein-free DNA replication junctions provide a structural framework for this working model. Those studies established that the junction rapidly interconverts among four representative conformational macrostates whose equilibrium populations depend on local DNA structure and solution conditions. In particular, transitions between the two dominant macrostates, S_2_ and S_3_, were proposed to occur through transient purine base flipping at the ss–dsDNA junction, accompanied by local rearrangements of base stacking and backbone geometry (3). Such transitions provide a plausible physical pathway by which the junction can rapidly explore multiple conformations while maintaining an overall intact DNA structure. Within this framework, the increased conformational disorder observed by 2DFS is consistent with CTD-mediated redistribution of equilibrium populations among conformational states already accessible to the protein-free junction, resulting in enhanced sampling of less-populated conformations within its intrinsic free-energy landscape.

Additional support for this interpretation comes from comparison of the 3′9A/5′2T and 3′2A/5′9T junctions, whose short ssDNA arms restrict binding to a single gp32 molecule and are expected to position the CTD differently relative to the junction (Supplementary Fig. S4c,d). In our previous kinetic studies, the polarity dependence of gp32 nucleation was interpreted in terms of a preferred binding orientation in which the CTD is positioned toward the junction on the 3′9A/5′2T construct but away from the junction on the oppositely oriented 3′2A/5′9T construct (6). The present spectroscopic measurements provide more direct experimental support for this model. For the 3′9A/5′2T junction, wild-type gp32 shifts the ensemble toward a right-handed average structure, whereas removal of the CTD shifts the ensemble toward a left-handed average structure. In contrast, for the 3′2A/5′9T junction, where the CTD is predicted to be oriented away from the junction, wild-type gp32 and gp32 I* produce similar shifts toward a left-handed average structure. Thus, these results support a model in which the ability of the CTD to alter the conformational landscape depends on its orientation relative to the ss–dsDNA junction, providing a structural basis for the architecture-dependent recognition inferred previously from gp32 nucleation kinetics.

### Alt text

Four-panel schematic model of how the gp32 CTD affects conformational ensembles of DNA replication junctions. Panel a shows four interconverting DNA conformational macrostates, S_1_–S_4_, with pathways connecting the states and a base-flipping intermediate between the S_2_ and S_3_ states. Panel b depicts protein-free junction architectures as different equilibrium distributions among the same four macrostates. Panel c shows a DNA fork junction bound by wild-type gp32 together with conformational distributions in which differences in macrostate populations among junction architectures are amplified. Panel d shows a DNA fork junction bound by CTD-truncated gp32 I* together with similar conformational distributions for different junction architectures.

The working model summarized in Fig. 6 also provides a broader perspective on how proteins achieve selective recognition of dynamic DNA substrates. Classical descriptions of protein access to duplex DNA generally invoke two limiting mechanisms. In the breathing-and-trapping model, proteins selectively capture transient conformations that arise through intrinsic thermal fluctuations of the DNA, whereas in the protein-induced opening model, the protein drives the DNA toward a conformation that is only weakly populated in the absence of protein (22,23). Although these two mechanisms provide useful conceptual frameworks, our results suggest that selective recognition by gp32 does not fit neatly into either category.

Rather than simply trapping rare pre-existing conformations or actively inducing a unique DNA structure, the intrinsically disordered CTD appears to couple dynamically to the intrinsic conformational landscape of the replication junction. We propose that transient CTD interactions redistribute the equilibrium populations among pre-existing conformational macrostates, thereby amplifying the architecture-dependent conformational disorder of the junction (Fig. 6). Selective molecular recognition may therefore arise not through stabilization of a single DNA conformation, but through amplification of differences among conformational distributions that already exist within the protein-free DNA junctions. More broadly, these findings suggest that intrinsically disordered protein domains can promote selective molecular recognition by reshaping pre-existing conformational landscapes of their nucleic acid substrates rather than by imposing a unique bound structure.

## Data Availability

The data supporting the findings of this study are deposited in Zenodo (24).

## Code Availability

Custom Python code used for Holstein–Frenkel modeling and analysis of the absorbance, circular dichroism, and two-dimensional fluorescence spectroscopy data is publicly available through the Marcus laboratory GitHub repository. The version of the code used in the current study is archived on Zenodo (24). The response-function database required for the 2DFS analysis is also archived on Zenodo (25).

## Supplementary Data

Supplementary Data are available at NAR Online.

## Acknowledgements

The authors thank Prof. Richard Karpel for providing the gp32 I* expression plasmid and Steve Weitzel and Larry Scatena for their support throughout this project. We also thank Prof. Marina Guenza and members of our laboratories for helpful discussions. P.H.v.H. is an American Cancer Society Research Professor of Chemistry.

## Author contributions

L.E., A.H.M. and P.H.v.H. conceived and designed the study. L.E. performed the experiments and analyzed the data. L.E. and A.H.M. contributed to experimental troubleshooting and interpretation of the results. L.E. and A.H.M. wrote the manuscript. P.H.v.H. contributed scientific consultation and critical review of the manuscript. All authors reviewed and approved the final manuscript.

## Funding

This work was supported by the National Institutes of Health, National Institute of General Medical Sciences [R35GM161360 to A.H.M.].

## Competing interests

The authors declare no competing interests.

## Notes

https://zenodo.org/records/22649211

https://zenodo.org/records/22102458

### Competing Interest Statement

The authors have declared no competing interest.

