## Supplementary Information for "The intrinsically disordered C-terminal domain of bacteriophage T4 gp32 amplifies pre-existing conformational heterogeneity at DNA replication junctions"

### Supplementary Materials and Methods

DNA constructs and protein preparation
Absorbance and circular dichroism spectroscopy
Two-dimensional fluorescence spectroscopy
Holstein–Frenkel modeling and spectral analysis

### Supplementary Tables

**Supplementary Table S1.** DNA construct sequences and nomenclature
**Supplementary Table S2.** Optimized structural and lineshape parameters

### Supplementary Figures

**Supplementary Fig. S1.** Representative complete optimization for the 3′9A/5′9T reference junction
**Supplementary Fig. S2.** Absorbance spectra of $(Cy3)_2$-labeled DNA replication junctions in the absence and presence of gp32
**Supplementary Fig. S3.** Circular dichroism spectra of $(Cy3)_2$-labeled DNA replication junctions in the absence and presence of gp32
**Supplementary Fig. S4.** Absorbance and circular dichroism spectra of DNA replication junctions in the absence and presence of wild-type gp32 and gp32 I*
**Supplementary Fig. S5.** Experimental and optimized two-dimensional fluorescence spectra for the DNA replication junction constructs and their gp32 complexes

**DNA constructs and protein preparation.** The $(Cy3)_2$-labeled ss–dsDNA replication junction constructs were assembled from pairs of complementary oligonucleotides synthesized by Integrated DNA Technologies (Coralville, IA). The constructs contained an internally linked Cy3 chromophore on each strand at the −1 position adjacent to the ss–dsDNA junction. The lengths of the 3′ and 5′ ssDNA arms were varied systematically while maintaining a common duplex region and probe position. An additional construct, 3′22T/5′9A*, was prepared in which A and T were exchanged between the two strands at the base pair immediately adjacent to the $(Cy3)_2$ probe. Complete oligonucleotide sequences and construct nomenclature are provided in Supplementary Table S1.

Lyophilized oligonucleotides were resuspended to 100 μM in 10 mM Tris and 0.1 mM EDTA (pH 8.0) and stored at −20 °C. Complementary strands were combined at a 1:1 molar ratio and annealed overnight using a programmable thermal controller. Unless otherwise indicated, all spectroscopic measurements were performed in 100 mM NaCl, 6 mM $MgCl_2$, and 10 mM Tris (pH 7.5).

Wild-type gp32 was prepared and purified as described previously (1). The gp32 I* expression plasmid was provided by Prof. Richard Karpel, and gp32 I* was expressed and purified using the same procedure. Purified proteins were dialyzed into the experimental buffer, and protein concentrations were determined spectrophotometrically at 280 nm using a molar extinction coefficient of $\varepsilon_{280}$ = 37,000 $M^{-1}$ $cm^{-1}$ for both wild-type gp32 and gp32 I* (2).

**Absorbance and circular dichroism spectroscopy.** Absorbance and circular dichroism (CD) spectra were measured using a JASCO J-720 spectropolarimeter with samples in 10-mm-path-length quartz cuvettes. Spectra were recorded from 400 to 650 nm with the sample maintained at 21 °C using a computer-controlled temperature stage. DNA concentrations were 1 μM for all absorbance and CD measurements.

Saturating stoichiometric concentrations of wild-type gp32 and gp32 I* were determined independently for each DNA construct by protein titration while monitoring changes in the CD spectrum. Protein was added until further additions produced no significant spectral change. Saturation was achieved at DNA:protein molar ratios of 1:3 for the 3′9A/5′2T and 3′2A/5′9T constructs, 1:4 for the 3′9A/5′9T reference construct, and 1:5 for the 3′22A/5′9T and 3′9A/5′22T constructs. Titrations were performed in duplicate. The 3′22T/5′9A* A/T-exchanged construct was measured only in the absence of protein.

**Two-dimensional fluorescence spectroscopy.** Phase-modulated two-dimensional fluorescence spectroscopy (2DFS) measurements were performed using the experimental approach described previously (3-5). Briefly, the output of a PHAROS laser (Light Conversion; 1030 nm, 250 kHz repetition rate) was used to pump a custom-built noncollinear optical parametric amplifier (NOPA), producing visible pulses centered at approximately 533 nm with a spectral bandwidth of approximately 33 nm. The pulses were compressed using a quadruple-pass fused-silica prism compressor, yielding a pulse-pair autocorrelation width of approximately 33 fs, corresponding to an estimated Gaussian pulse duration of approximately 23 fs. The excitation spectrum was monitored throughout each measurement.

Samples were contained in a 1-mm-path-length quartz cuvette and continuously circulated using a peristaltic pump at a flow rate of approximately 85 μL $min^{-1}$ to minimize optical saturation and photodegradation. Fluorescence was detected with a photomultiplier tube (Hamamatsu R1527) following a 615-nm long-pass filter to suppress scattered excitation light.

The population time between the second and third excitation pulses was set to $T = 0$ for all measurements. Rephasing (RP) and nonrephasing (NRP) signals were acquired as functions of the coherence time delays and Fourier transformed to obtain the corresponding two-dimensional frequency-domain spectra. The frequency axes were referenced using optical reference signals as described previously (3).

At the beginning of each experimental session, measurements of the +15 duplex DNA reference sample described previously (4) were used to verify instrument performance and measurement reproducibility. Photomultiplier-tube and lock-in-amplifier settings were optimized using the reference sample and subsequently held constant for all Cy3-labeled samples and across experimental sessions. Before each measurement, the excitation spectrum, optical power, and temporal overlap of the excitation pulse sequence were monitored, and the temporal overlap was recalibrated as necessary to compensate for drift in the delay-stage positions.

For each DNA construct, two measurements were initially acquired in the absence of protein to assess reproducibility. The sample was then removed from the cuvette using the peristaltic pump, and protein was added at the appropriate saturating stoichiometric concentration. The protein–DNA complex was incubated for 10 min at room temperature before being returned to the cuvette for measurement. This procedure was repeated for measurements at different protein concentrations or with different protein–DNA complexes.

Between samples, the cuvette and sample-flow system were thoroughly cleaned to minimize sample carryover. The NOPA output spectrum and optical power were monitored throughout data acquisition, and the instrument was recalibrated when necessary. Following recalibration, measurements of the reference DNA sample were repeated to confirm instrument performance before additional experimental samples were measured.

**Holstein–Frenkel modeling and spectral analysis.** Absorbance and CD spectra of the $(Cy3)_2$-labeled DNA junctions were modeled using the Holstein–Frenkel Hamiltonian for a coupled chromophore dimer, as described previously (4-6). Excitonic coupling between the Cy3 chromophores splits the vibronic transitions into symmetric and antisymmetric exciton states whose energies and rotational strengths depend sensitively on the relative positions and orientations of the two chromophores.

The electronic coupling was calculated using a transition-charge description in which transition electrostatic potential (TrESP) charges are assigned to the atomic coordinates of each Cy3 chromophore (6,7). The relative Cy3 dimer geometry was parameterized by three angular coordinates $\phi$, $\theta$, and $\eta$) and three translational coordinates $R$, $\delta$, and $\xi$). These parameters were determined by simultaneous numerical optimization of the calculated absorbance and CD spectra against the corresponding experimental spectra, as described previously (6). Representative fits for the 3′9A/5′9T reference construct are shown in Supplementary Fig. S1, and optimized absorbance and CD spectra for the complete construct series are shown in Supplementary Fig. S4.

The same Holstein–Frenkel Hamiltonian was subsequently used to calculate the RP and NRP 2DFS response functions as described previously (4,8,9). The calculated fluorescence-detected response includes contributions from ground-state bleach (GSB), stimulated emission (SE), and excited-state absorption (ESA) pathways. The relative contribution of the doubly excited-state manifold is governed by the parameter $\Gamma_{2D}$, which accounts for the fluorescence quantum yield of the doubly excited states relative to that of the singly excited states. This parameter was fixed at 0.3, as determined previously for the $(Cy3)_2$ probe (4). Finite excitation bandwidth and rotational averaging for the parallel-polarization pulse sequence were incorporated as described previously (9).

For each sample, the structural parameters obtained from the simultaneous absorbance/CD optimization were held fixed during the analysis of the 2DFS data. The homogeneous and inhomogeneous linewidth parameters, $\Gamma_H$ and $\sigma_I$, respectively, were then determined by simultaneous optimization of the RP and NRP signals. The differing dependence of the RP and NRP lineshapes on homogeneous and inhomogeneous broadening permits these contributions to be distinguished. Optimized frequency-domain spectra were generated by Fourier transformation of the calculated RP and NRP response functions and compared directly with experiment (Supplementary Figs. S1 and S5). Parameter uncertainties were determined following the procedure described previously (4), with uncertainty bounds defined for the present analysis by parameter variations satisfying $\Delta\chi^2/\chi_0^2 = 0.005$, corresponding to a 0.5% increase relative to the optimized $\chi_0^2$ value. The resulting structural and lineshape parameters are summarized in Supplementary Table S2.

**Supplementary Table S1. Base sequences and nomenclature of the exciton-coupled $(Cy3)_2$-labeled DNA constructs used in this study.** In all constructs, the $(Cy3)_2$ exciton probe was positioned at the −1 site immediately adjacent to the ss–dsDNA junction (4). The lengths of the 3′ and 5′ ssDNA arms were varied while maintaining a common duplex region and probe position. Construct names specify the lengths (nt) of the 3′ and 5′ ssDNA arms. The suffix "A" or "T" specifies whether the corresponding ssDNA arm is contiguous with the adenine- or thymine-containing strand, respectively, of the terminal A:T base pair adjacent to the $(Cy3)_2$ probe.

| Construct | Strand 1 (5' → 3') \| Strand 2 (3' → 5') |
|---|---|
| 3'9A / 5'2T | 5'–CTCCCTCGTGTCGTCTCCAGTCATAATATGCGA(Cy3)ATACTTTCG–3'<br>3'–GAGGGAGCACAGCAGAGGTCAGTATTATACGCT(Cy3)CG–5' |
| 3'22A / 5'9T | 5'–CTCCCTCGTGTCGTCTCCAGTCATAATATGCGA(Cy3)ATACTTTCG**TTTTTTTTTTTTT**–3'<br>3'–GAGGGAGCACAGCAGAGGTCAGTATTATACGCT(Cy3)CGCTGGTAT–5' |
| 3'9A / 5'9T | 5'–CTCCCTCGTGTCGTCTCCAGTCATAATATGCGA(Cy3)ATACTTTCG–3'<br>3'–GAGGGAGCACAGCAGAGGTCAGTATTATACGCT(Cy3)CGCTGGTAT–5' |
| 3'9A / 5'22T | 5'–CTCCCTCGTGTCGTCTCCAGTCATAATATGCGA(Cy3)ATACTTTCG–3'<br>3'–GAGGGAGCACAGCAGAGGTCAGTATTATACGCT(Cy3)CGCTGGTAT**TTTTTTTTTTTTT**–5' |
| 3'2A / 5'9T | 5'–CTCCCTCGTGTCGTCTCCAGTCATAATATGCGA(Cy3)AT–3'<br>3'–GAGGGAGCACAGCAGAGGTCAGTATTATACGCT(Cy3)CGCTGGTAT–5' |
| 3'22T / 5'9A* | 5'–CTCCCTCGTGTCGTCTCCAGTCATAATATGCGT(Cy3)ATACTTTCG**TTTTTTTTTTTTT**–3'<br>3'–GAGGGAGCACAGCAGAGGTCAGTATTATACGCA(Cy3)CGCTGGTAT–5' |

* The 3′22T/5′9A* construct has the same junction architecture as the 3′22A/5′9T construct, but with A and T exchanged between the two strands at the terminal base pair immediately adjacent to the $(Cy3)_2$ probe.

**Supplementary Table S2.** Optimized structural and lineshape parameters obtained from global analysis of the absorbance, circular dichroism (CD), and two-dimensional fluorescence spectroscopy (2DFS) measurements. Ensemble-averaged structural parameters of the $(Cy3)_2$ probe were determined by simultaneous optimization of the absorbance and CD spectra using a Holstein-Frenkel Hamiltonian in combination with the transition-charge model (6). The homogeneous and inhomogeneous linewidth parameters, $\Gamma_H$ and $\sigma_I$, respectively, were subsequently obtained by simultaneous optimization of the RP and NRP 2DFS spectra while holding the structural parameters fixed. Reported uncertainties were determined from the fitting procedure using criteria $\Delta\chi^2/\chi_0^2$ = 0.005, corresponding to a 0.5% increase relative to the optimized $\chi_0^2$ value.

| Con-struct | Com-plex | Structural parameters from Holstein-Frenkel analysis of the absorbance and CD spectra | | | | | | | Lineshape parameters from 2DFS analysis | |
|---|---|---|---|---|---|---|---|---|---|---|
| | | $\phi$ (º) | $\theta$ (º) | $\eta$ (º) | $\xi$ (Å) | $\delta$ (Å) | $R$ (Å) | $J$ (cm$^{-1}$) | $\Gamma_H$ (cm$^{-1}$) | **$\sigma_I$ (cm$^{-1}$)** |
| 3’9A / 5’2T | DNA | 73 ±1 | 22 ±2 | 112 ±7 | -3.5 ±1 | 0.45 ±1 | 9.0 ±0.2 | 351 ±10 | 155 ± 6 | **204 ±10** |
| | WT gp32 | 75 ±1 | 29 ±2 | 93 ±10 | -2.72 ±1 | -1.8 ±1 | 8.4 ±0.2 | 346 ±10 | 204 ± 7 | **211 ±11** |
| | gp32 I* | 102 ±1 | 11 ±10 | 114 ±3 | -0.14 ±1 | -3.15 ±1 | 7.1 ±0.2 | -374 ±10 | 170 ± 7 | **206 ±10** |
| 3’22A / 5’9T | DNA | 103 ±1 | 13 ±6 | 124 ±3 | 0.86 ±1 | 4.5 ±1 | 8.4 ±0.4 | -384 ±10 | 162 ± 8 | **205 ±13** |
| | WT gp32 | 75 ±1 | 3.0 ±1 | 96 ±1 | -0.54 ±1 | -4.7 ±1 | 8.0 ±0.2 | 392 ±10 | 175 ± 9 | **220 ±11** |
| | gp32 I* | 92 ±1 | 29 ±1 | 116 ±1 | 0.8 ±1 | -3.9 ±1 | 6.5 ±0.2 | 430 ±10 | 187 ± 8 | **191 ±11** |
| 3’9A / 5’9T | DNA | 99 ±1 | 40 ±10 | 118 ±5 | 1.30 ±1 | 3.72 ±1 | 7.4 ±1 | -402 ±25 | 160 ± 5 | **180 ± 9** |
| | WT gp32 | 69 ±1 | 15 ±1 | 125 ±3 | -3.67 ±1 | -0.4 ±1 | 8.5 ±0.2 | 466 ±15 | 160 ± 6 | **197 ±10** |
| | gp32 I* | 94 ±1 | 16 ±3 | 113 ±1 | 0.38 ±1 | 3.72 ±1 | 6.3 ±0.2 | -426 ±15 | 173 ± 8 | **192 ±12** |
| 3’9A / 5’22T | DNA | 89.8 ±1 | 29 ±1 | 123 ±1 | 0.75 ±1 | -4.75 ±1 | 7.3 ±0.2 | -486 ±15 | 214 ± 10 | **169 ±15** |
| | WT gp32 | 73 ±1 | 28 ±1 | 108 ±5 | -3.47 ±1 | -0.31 ±1 | 8.7 ±0.2 | 439 ±15 | 184 ± 6 | **182 ±10** |
| | gp32 I* | 97 ±1 | 21±5 | 120 ±3 | 0.49 ±1 | -3.6 ±1 | 7.0 ±0.2 | -397 ±15 | 179 ± 8 | **191 ±12** |
| 3’2A / 5’9T | DNA | 71 ±1 | 24 ±2 | 103 ±3 | -3.8 ±1 | 1.13 ±1 | 9.3 ±0.2 | 381 ±10 | 205 ± 7 | **192 ±11** |
| | WT gp32 | 102 ±1 | 1.8 ±10 | 103 ±3 | -0.58 ±1 | 3.01 ±1 | 6.5 ±0.2 | -386 ±10 | 171 ± 7 | **224 ±12** |
| | gp32 I* | 102 ±1 | 19 ±2 | 121 ±1 | -0.07 ±1 | -2.6 ±1 | 6.9 ±0.2 | -389 ±10 | 181 ± 6 | **203 ±11** |
| 3’22T / 5’9A* | DNA | 67 ±1 | 4.8 ±1 | 112 ±3 | 0.63 ±1 | 2.6 ±1 | 6.8 ±0.2 | 437 ±15 | 171 ± 6 | **174 ± 9** |

* The 3′22T/5′9A* construct has the same junction architecture as the 3′22A/5′9T construct, but with A and T exchanged between the two strands at the terminal base pair immediately adjacent to the $(Cy3)_2$ probe.

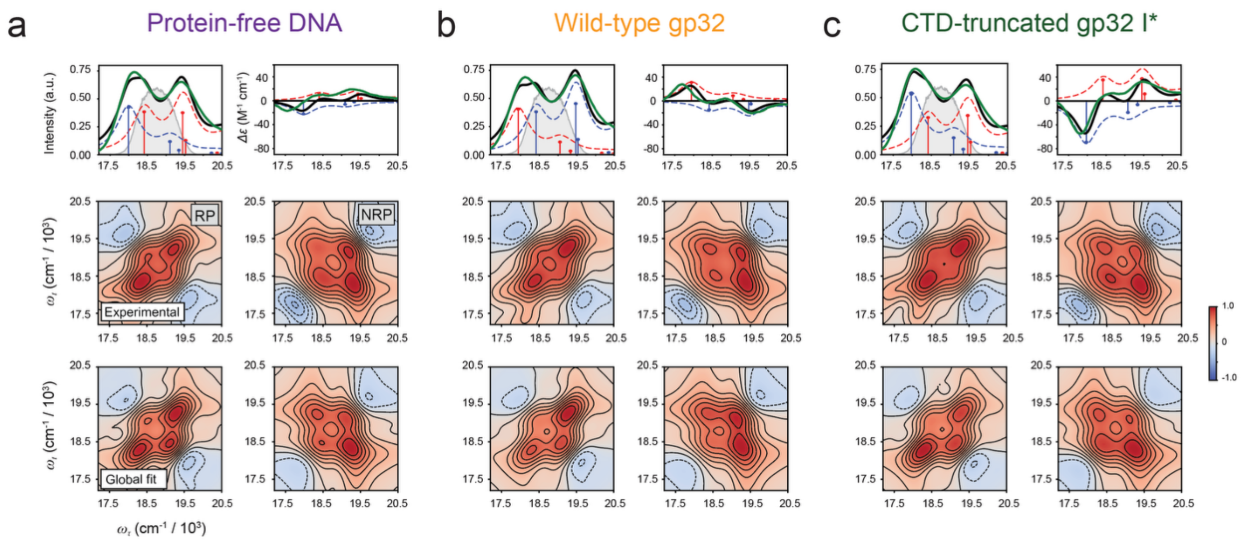


**Supplementary Fig. S1. Optimization of the absorbance, circular dichroism (CD), and two-dimensional fluorescence spectroscopy (2DFS) spectra for the 3′9A/5′9T DNA replication junction.** **(a)** Protein-free DNA. **(b)**Wild-type gp32–DNA complex. **(c)** gp32 I*–DNA complex. The upper row shows the experimental absorbance and CD spectra together with the optimized calculated spectra. Black curves represent the experimental spectra and green curves the optimized calculated spectra; red and blue curves denote the individual vibronic components. The absorbance panels also show the laser excitation spectrum (gray). The middle and lower rows compare the experimental and optimized rephasing (RP) and nonrephasing (NRP) 2DFS spectra, respectively. Structural parameters were determined by simultaneous optimization of the absorbance and CD spectra, after which the homogeneous and inhomogeneous linewidth parameters, $\Gamma_H$ and $\sigma_I$, were determined by simultaneous optimization of the RP and NRP 2DFS spectra while holding the structural parameters fixed. The optimized model provides a consistent description of the absorbance, CD, RP, and NRP spectra. Optimized parameters are listed in Supplementary Table S2.

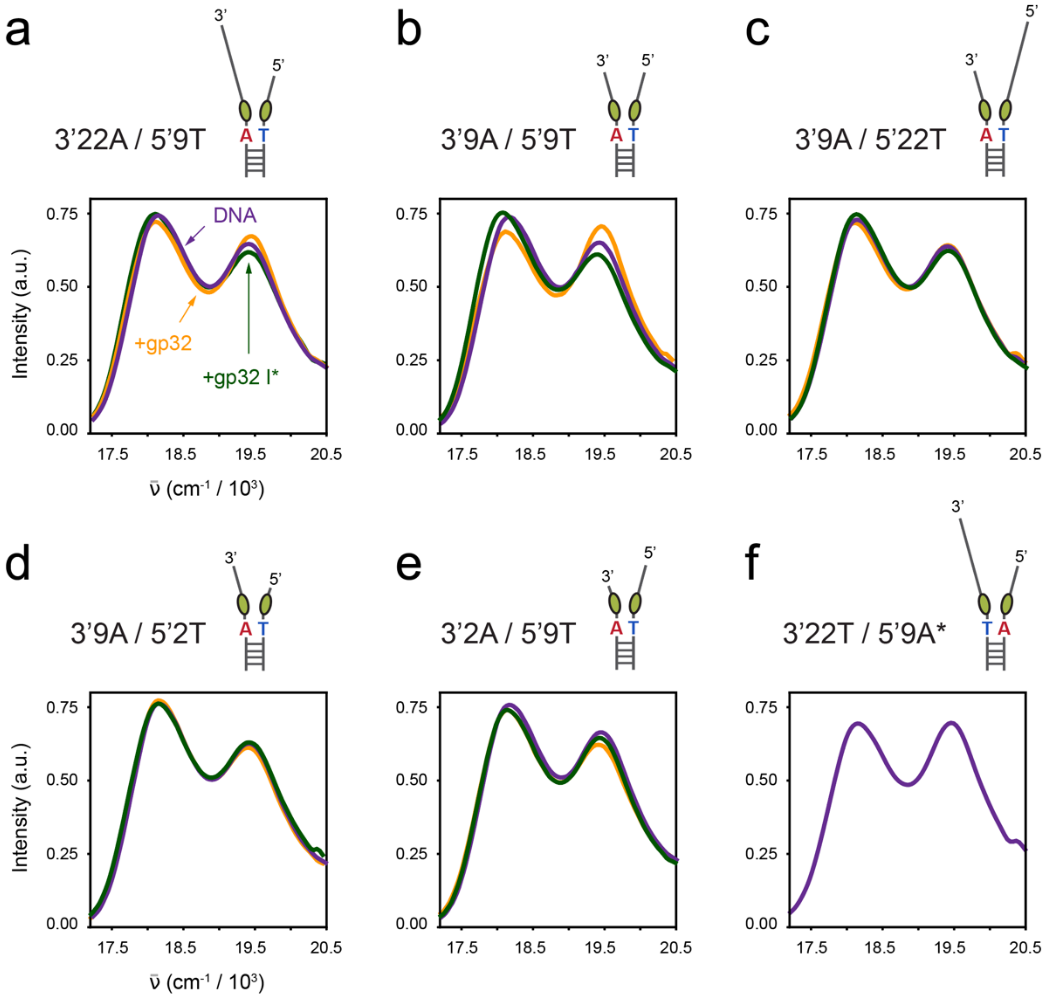


**Supplementary Fig. S2. Absorbance spectra of $(Cy3)_2$-labeled DNA replication junctions in the absence and presence of gp32.** Absorbance spectra normalized to equal integrated area are shown for the six DNA junction constructs: **(a)** 3′22A/5′9T, **(b)** 3′9A/5′9T, **(c)** 3′9A/5′22T, **(d)** 3′9A/5′2T, **(e)** 3′2A/5′9T, and **(f)** 3′22T/5′9A*. Spectra are shown for protein-free DNA (purple), complexes with wild-type gp32 (orange), and complexes with CTD-truncated gp32 I* (green). For the 3′22T/5′9A* A/T-exchanged control **(f)**, measurements were performed only for protein-free DNA. Junction cartoons above each panel indicate the lengths of the 3′ and 5′ ssDNA arms and the orientation of the terminal A:T base pair adjacent to the $(Cy3)_2$ probe.

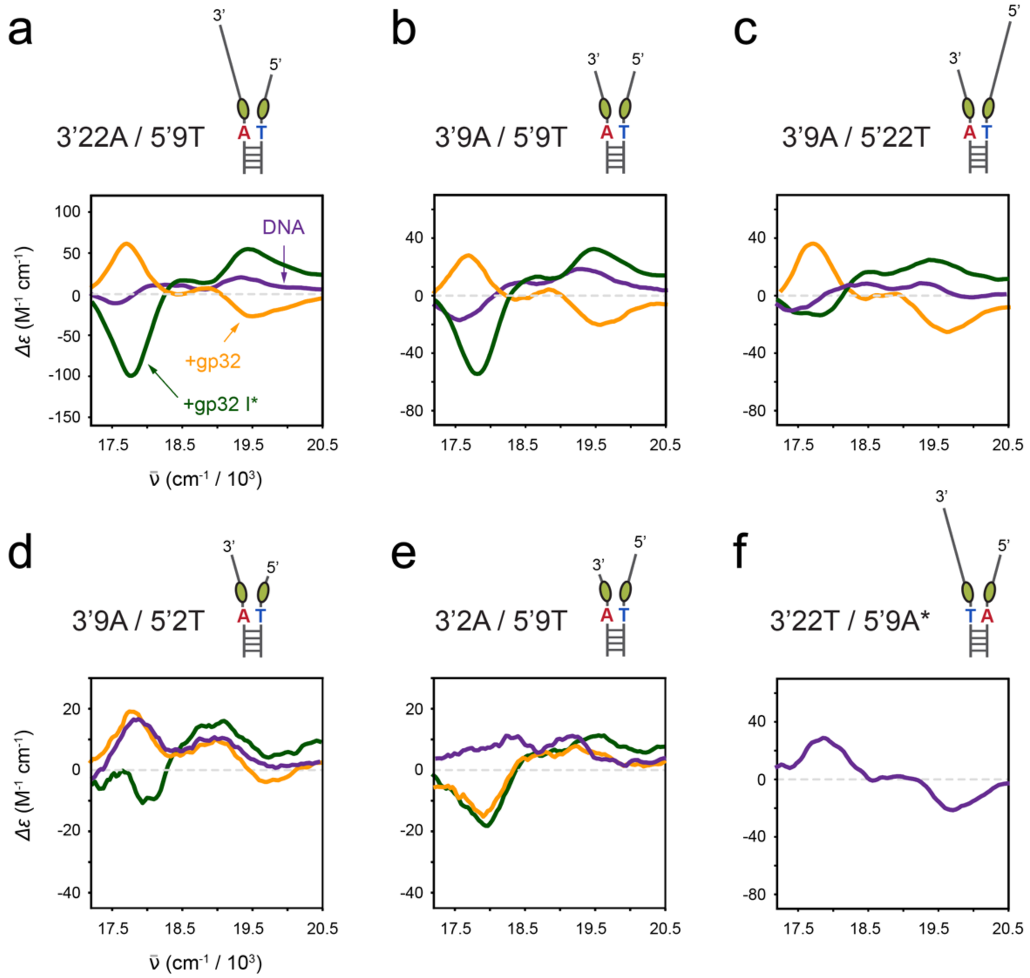


**Supplementary Fig. S3. Circular dichroism spectra of $(Cy3)_2$-labeled DNA replication junctions in the absence and presence of gp32.** CD spectra are shown for the six DNA junction constructs: **(a)** 3′22A/5′9T, **(b)** 3′9A/5′9T, **(c)**3′9A/5′22T, **(d)** 3′9A/5′2T, **(e)** 3′2A/5′9T, and **(f)** 3′22T/5′9A*. Spectra are shown for protein-free DNA (purple), complexes with wild-type gp32 (orange), and complexes with CTD-truncated gp32 I* (green). For the 3′22T/5′9A* A/T-exchanged control **(f)**, measurements were performed only for protein-free DNA. Junction cartoons above each panel indicate the lengths of the 3′ and 5′ ssDNA arms and the orientation of the terminal A:T base pair adjacent to the $(Cy3)_2$ probe. Note that the ordinate scales differ among panels to accommodate the substantial differences in CD amplitude across junction architectures.

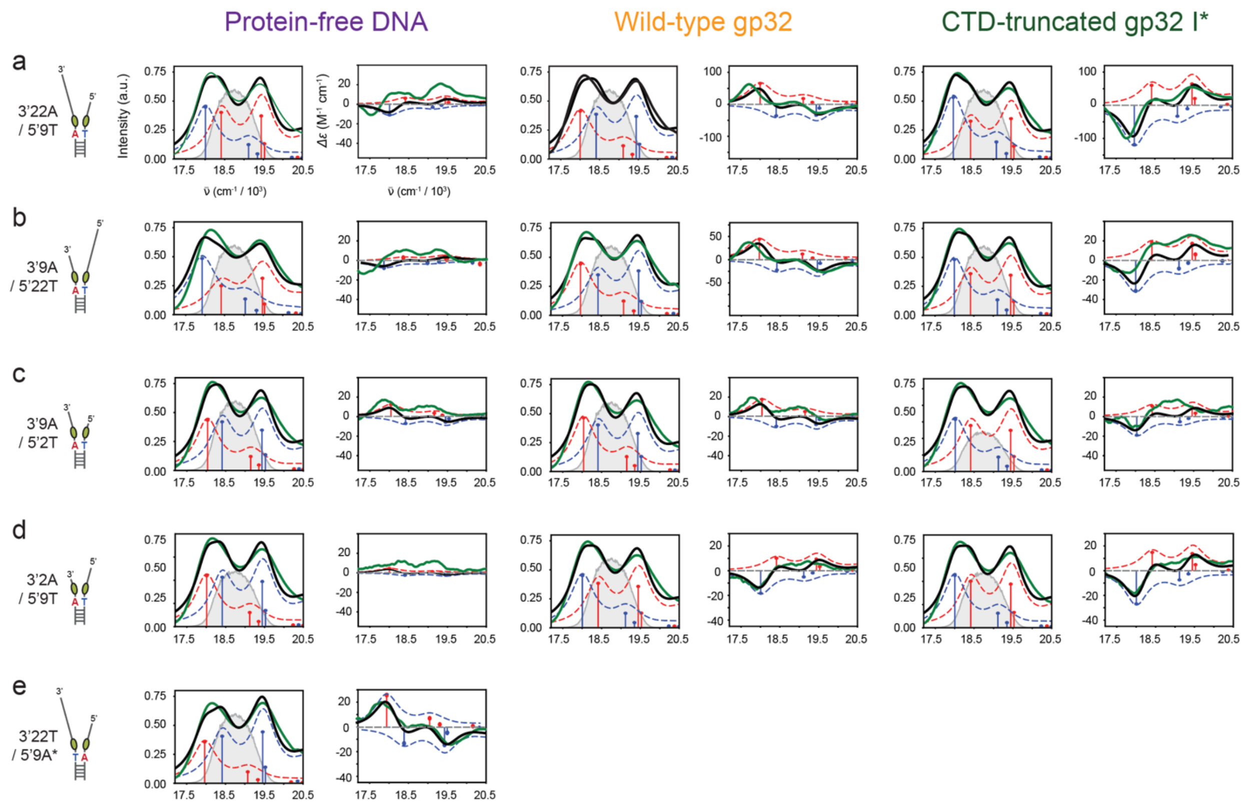


**Supplementary Fig. S4. Absorbance and circular dichroism spectra of DNA replication junctions in the absence and presence of wild-type gp32 and gp32 I*.** Experimental absorbance and circular dichroism (CD) spectra are shown for the indicated DNA replication junction constructs as protein-free DNA (left), complexes with wild-type gp32 (middle), and complexes with the CTD-truncated gp32 I* mutant (right). Black curves represent the experimental spectra and green curves the optimized spectra calculated using the Holstein–Frenkel model; red and blue curves denote the individual vibronic components. The laser excitation spectrum is shown in gray in the absorbance panels and indicates the spectral region sampled in the corresponding 2DFS measurements shown in Supplementary Fig. S5. **(a)** 3′22A/5′9T. **(b)**3′9A/5′22T. **(c)** 3′9A/5′2T. **(d)** 3′2A/5′9T. **(e)** 3′22T/5′9A*, for which protein-complex spectra were not measured. The single-gp32-binding constructs in **(c)** and **(d)** provide a comparison of complexes in which the CTD is predicted to be oriented differently relative to the ss–dsDNA junction based on previous gp32 nucleation measurements (10). Optimized structural parameters are listed in Supplementary Table S2.

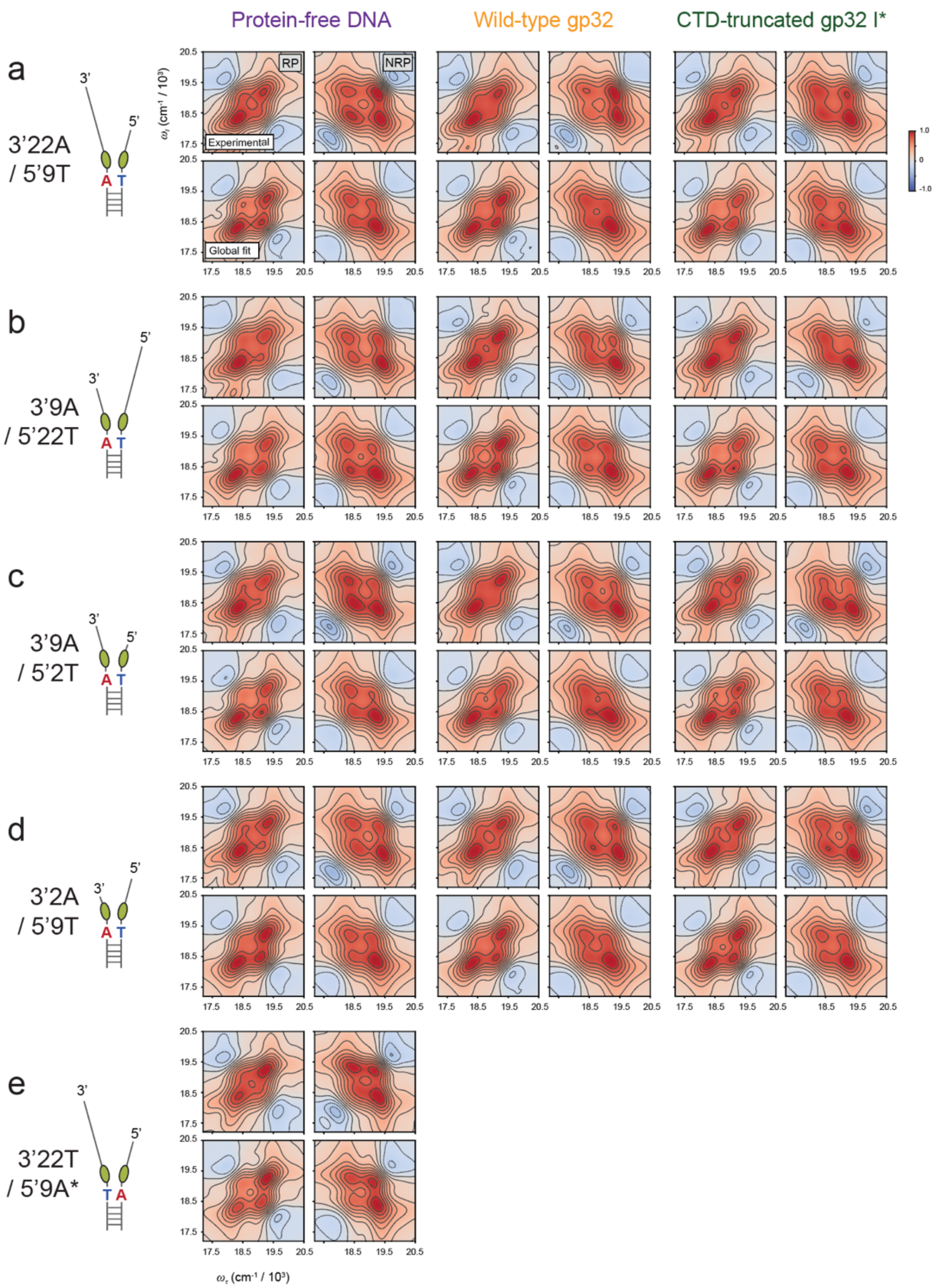


**Supplementary Fig. S5. Experimental and optimized two-dimensional fluorescence spectra for the DNA replication junction constructs and their gp32 complexes. (a–e)** Rephasing (RP) and nonrephasing (NRP) 2DFS spectra are shown for the indicated DNA junction constructs. For each condition, the upper row shows the experimental spectra, and the lower row shows the corresponding spectra calculated using the optimized model parameters. Columns compare protein-free DNA, wild-type gp32–DNA complexes, and gp32 I*–DNA complexes. Structural parameters were fixed at values obtained from simultaneous optimization of the absorbance and CD spectra, and the homogeneous and inhomogeneous linewidth parameters, $\Gamma_H$ and $\sigma_I$, were determined by simultaneous optimization of the corresponding RP and NRP 2DFS spectra. No protein-complex measurements were obtained for the 3′22T/5′9A* construct. Optimized parameters are listed in Supplementary Table S2.
